# A timeline for evolution of metabolism in early microbes

**DOI:** 10.64898/2026.11.02.763483

**Authors:** Timothy J. Hackmann

## Abstract

The metabolism of early microbes shaped our planet’s chemistry and enabled more life to evolve. However, it remains uncertain when many forms of metabolism arose and which organisms carried them out. Here we combine machine learning, metabolic networks, and phylogenomics of microbes to date 14 forms of metabolism and 6 other traits. We find that metabolism evolved in three waves, starting >3.4 billion years ago (Ga) with fermentation and sulfur compound respiration. A second wave began ∼3.2 Ga with aerobic respiration and growth, while a third began ∼1.4 Ga with further specialization. We compared these predictions with independent dates from the geochemical record, and agreement with conservative dates was good (CCC = 0.81 to 0.89). Several forms of metabolism are invisible in the geochemical record, and our approach dates these forms, too. Our results reconcile different lines of evidence for microbial evolution and provide a fuller picture of early Earth.

## INTRODUCTION

Early microbes shaped our planet through their metabolism (*1–3*). The first microbes to carry out photosynthesis, for example, released oxygen into the atmosphere and changed our planet’s chemistry (*4*). Microbes that carried out nitrogen fixation released organic nitrogen (*2*, *5*), paving the way for more life to evolve. Modern forms of metabolism are responsible for cycling of several elements in the environment (*6*, *7*). They are also responsible for converting light and chemical energy into forms usable by other organisms (*6*, *7*).

Despite the importance, it has been hard to identify when major forms of metabolism arose and which organisms carried them out. The geochemical and fossil records are among the strongest means for identification (*1–3*), but they have ambiguities. Based on the geochemical record, photosynthesis has been estimated to have arisen anywhere from 3.2 to 2.35 billion years ago (Ga) (*4*, *8–11*). Some estimates put it even earlier (3.8 Ga) (*12*). The range is broad in other cases, such as with methanogenesis (3.5 to 2.7 Ga) (*13–15*). The fossil record can tie specific organisms (cyanobacteria) to photosynthesis (*16–18*). However, many forms of metabolism (e.g., fermentation) leave no trace in the fossil record or the geochemical one.

Genomics and computational methods may help fill gaps (Fig. S1). One method, gene reconstruction, uses genomes of modern organisms to predict genes of early ones (*19–21*). This approach has been applied to date genes or pathways for the sulfur cycle (*22*), nitrogen cycle (*23*, *24*), photosynthesis (*25*), and other forms of metabolism (*26*). A second method, trait prediction, is downstream of gene reconstruction. It uses genes and a computational algorithm to predict if whole traits are present or absent. In one study (*25*), a machine learning algorithm was trained on modern organisms to predict presence of aerobic growth. It was then applied to genes of early microbes, predicting that aerobic growth appeared at 3.2 Ga (*25*). Recent advances in trait prediction (*27*, *28*) could be used to date a range of traits, including metabolic ones, and build on dates for genes or pathways.

One challenge for genomics and computational methods is testing their predictions. Studies using these methods usually compare predicted dates to ones in the geochemical record (*22–26*). Because dates in the geochemical record are few, studies are often forced to use only one or two dates, or they use dates already used in time calibration. If more independent dates could be tested, it would provide even more support for these methods.

Here we use advances in genomics and computational methods to build a more complete and testable timeline for the evolution of metabolism. We reconstructed >9,300 gene functions of 700 microbes and their ancestors, then used two computational tools to predict the appearance of 14 forms of metabolism and 6 other traits. The dates they predicted agreed with independent dates in the geochemical record. Our results suggest that metabolism evolved in three waves, reconcile different lines of evidence for microbial evolution, and provide a more complete view of our early planet.

## RESULTS

### Our analysis used 700 genomes of bacteria and archaea

To build a timeline for metabolism, we needed to obtain a dated phylogenetic tree of organisms and their gene functions. We started with a set of 700 genomes of modern bacteria and archaea from Moody et al. (*26*, *29*, *30*). Those studies were also the source of an undated tree and gene functions of these modern organisms.

We dated the tree using fossil calibrations (Fig. 1A,B), choosing them to be as independent from the geochemical record as possible. This involved 10 nodes of the tree bounded by dates for 9 fossils and 1 geological event (Supplementary Text and Table S1). We excluded geochemical events, such as appearance of oxygen, so predictions from the tree could be independently compared to the geochemical record. The resulting tree showed that bacteria and archaea diverged from the last universal common ancestor (LUCA) at 4.48 Ga. The crown group for bacteria arose at 3.71 Ga, the group for archaea arose at 3.46 Ga, and they both diversified thereafter.

**Fig. 1.**
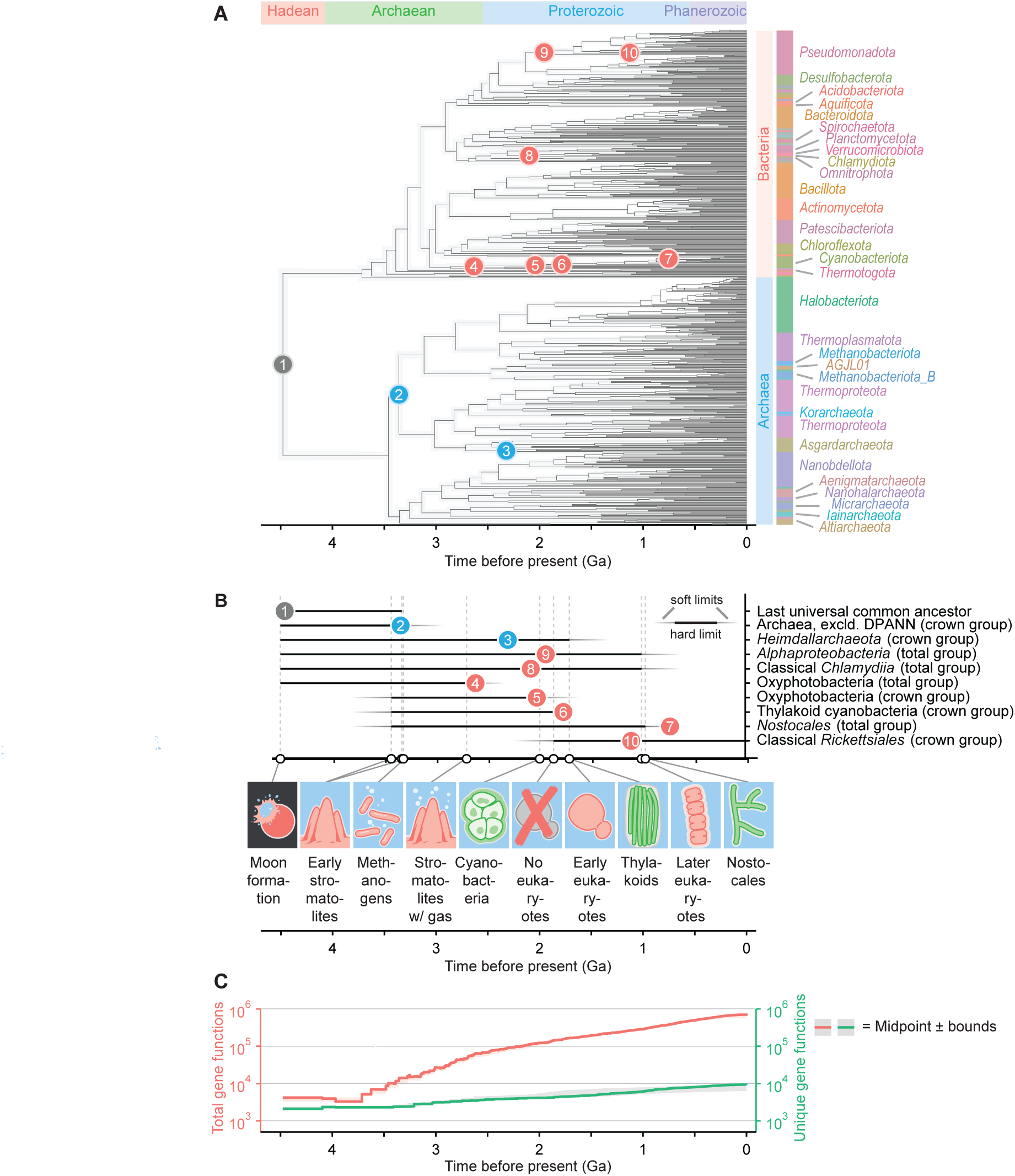
Our analysis used a phylogenetic tree of 700 genomes dated with fossils and 9,371 reconstructed gene functions. **(A)** Phylogenetic tree. **(B)** Fossils used for calibrations and bounds on tree nodes. **(C)** Reconstructed gene functions. Abbreviations: excld. = excluding and Ga = billion years ago.

We next reconstructed gene functions for the organisms in the tree (Fig. 1C). We used gene functions for modern organisms, the Count software package (*19*), and its gain-loss-duplication (GLD) model to perform the reconstruction. With this approach, we reconstructed 9,371 unique gene functions and 704,158 total gene functions (which included duplicates across lineages). The number started low and rose over time, showing diversification over evolutionary history. With the gene reconstruction and dated tree, we could proceed with predicting traits, including metabolism, over evolutionary time.

### Two tools predict the rise of major forms of metabolism

We input reconstructed genes into Microbe Decoder (*27*), a web server for predicting forms of metabolism and other traits. We predicted traits using two separate tools. The machine learning tool predicts if an organism has a trait using its gene functions and a random forest classifier. The classifier is trained on a dataset of modern organisms observed to have the trait. The metabolic networks tool predicts if an organism has a given metabolic pathway from its gene functions and a reference network. The reference network corresponds to all known reactions for a given type of metabolism, and gene functions are mapped to it. This tool originally handled only fermentative metabolism (*28*) but has been recently updated (*27*).

Our tools predicted the rise of 14 forms of metabolism (Data S1 and S2). Plotting predictions on trees (Fig. 2A) showed forms of metabolism that arose early and were widespread (e.g. fermentation and sulfur compound respiration). These plots also showed forms that arose late and remained more concentrated (e.g., selenate respiration and chlorate respiration). This pattern was also apparent when plotting the abundance of positive lineages (Fig. 2B and Fig. S2) and dating the appearance of the trait (first positive lineage) (Table S2).

**Fig. 2.**
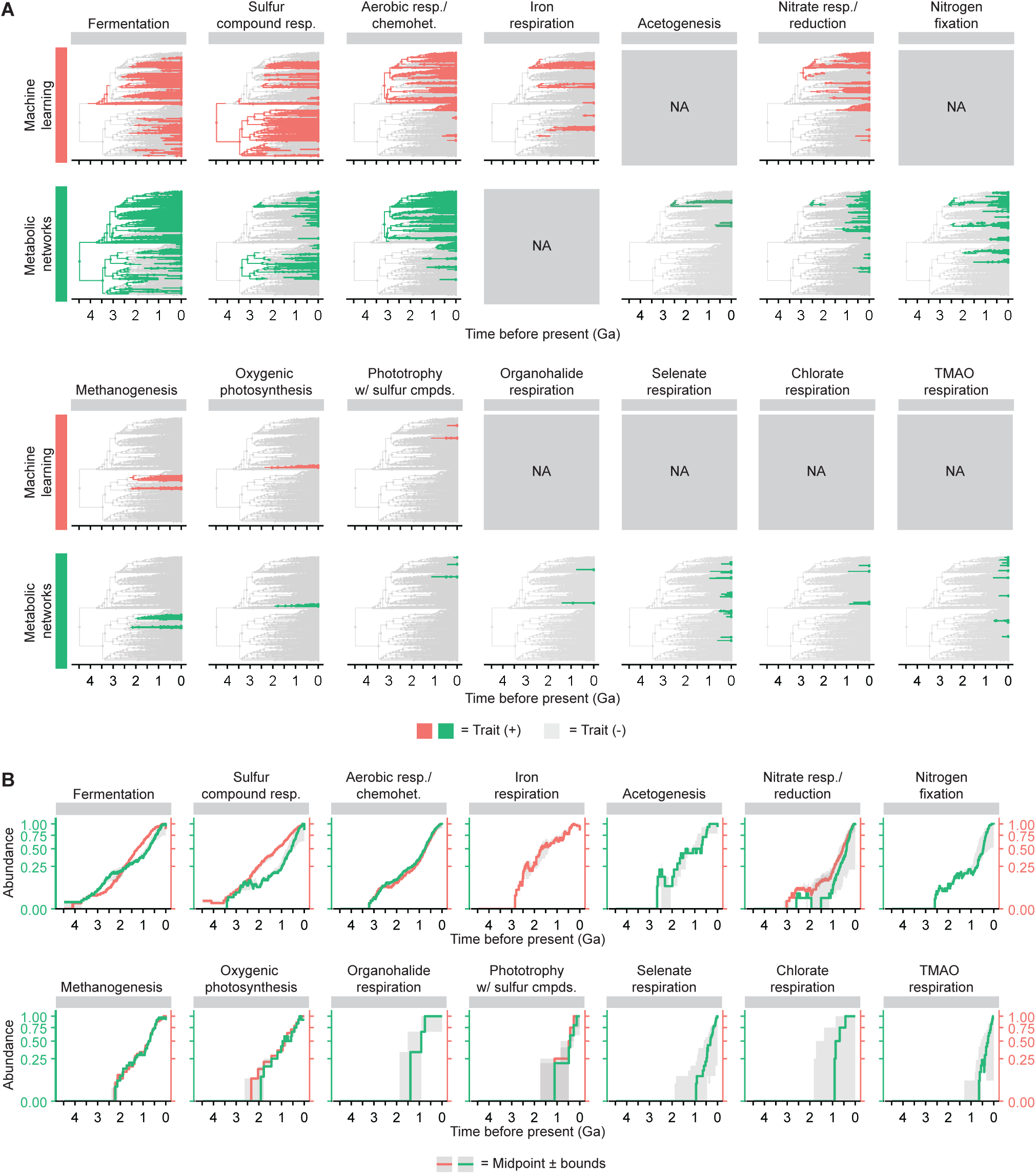
Our tools predict the rise of major forms of metabolism in microbes over evolutionary time. **(A)** Phylogenetic trees colored by predictions. **(B)** Abundance of lineages positive for a trait. In (B), values are scaled to the modern value and calculated as (lineages positive for trait x at time t) / (lineages positive for trait x at time 0). A value of 0.5 at 1 Ga means the number of lineages positive for the trait at that time was 50% of the number positive today. See Fig. S2 for values scaled by the historic value. Abbreviations: chemohet. = chemoheterotrophy, cmpds. = compounds, Ga = billion years ago, NA = prediction not available in Microbe Decoder, resp. = respiration, TMAO = trimethylamine N-oxide, and w/ = with.

Some forms of metabolism have a well-known taxonomic range, and our phylogenetic trees generally reproduce this pattern (Fig. 2A). For example, oxygenic photosynthesis was restricted to photosynthetic cyanobacteria (oxyphotobacteria [class *Cyanobacteriia*]). Additionally, methanogenesis was restricted to archaea (mostly phyla *Halobacteriota*, *Thermoplasmatota*, and *Methanobacteriota*). This supports the accuracy of the tools, which had been previously evaluated with modern microbes (*27*, *28*).

When a type of metabolism was predicted by both tools, the predictions generally agreed with each other (Fig. 2A,B and Fig. S3 and Table S2). The dates for appearance of traits, for example, were highly correlated between tools (*r* = 0.92 and CCC = 0.89) (Fig. S3). These two sets of predictions are helpful for building a timeline of evolution of metabolism.

### One tool predicts the rise of other traits

In addition to predicting forms of metabolism, we predicted 6 non-metabolic traits (Data S1). These are predicted by the machine learning tool of Microbe Decoder (*27*). These predictions showed that anaerobic, gram negative, and motile microbes arose early (Fig. 3). Aerobic, gram positive, and spore forming microbes arose later (Fig. 3). Aerobic growth arose at nearly the same time as aerobic respiration, a type of metabolism (Figs. 2 and 3). The agreement is notable given that the random forest classifiers for these two traits were trained on different datasets (*27*). Thus, results for these traits complement those for metabolism and provide a fuller picture of microbes over time.

**Fig. 3.**
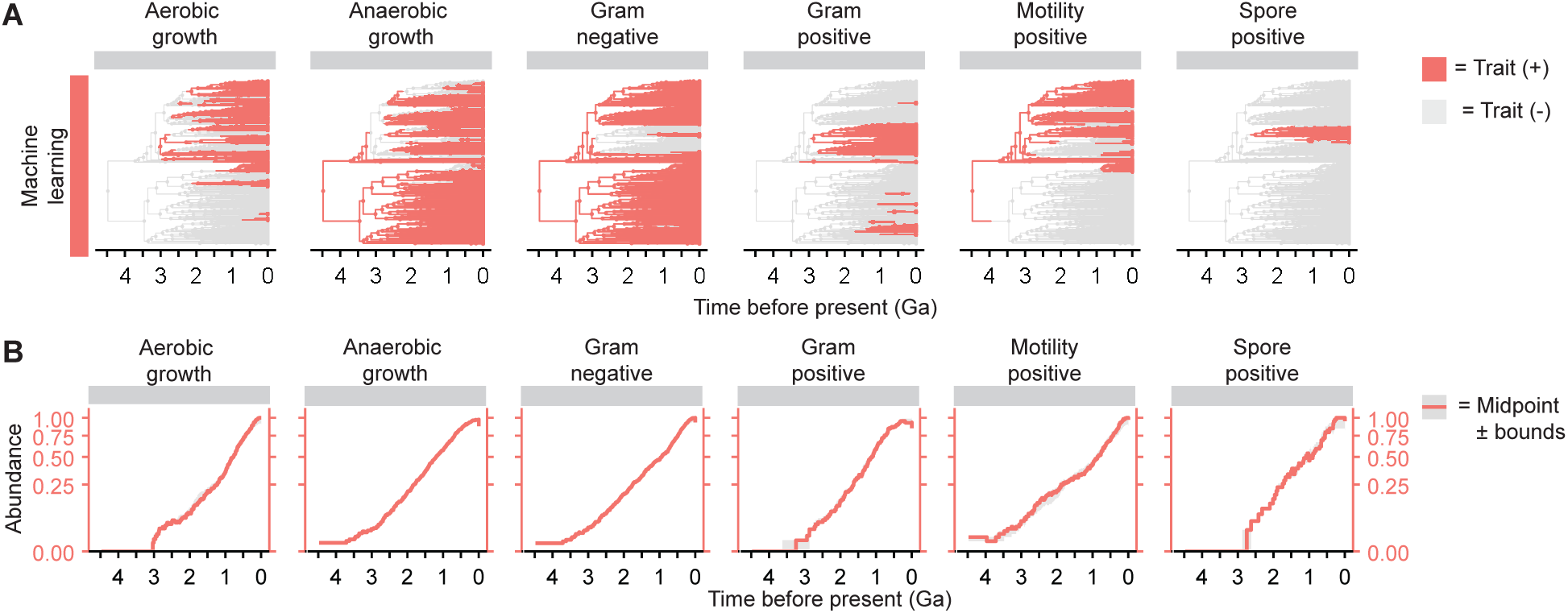
Our tools predict the rise of aerobic metabolism and other traits in microbes over evolutionary time. **(A)** Phylogenetic trees colored by predictions. **(B)** Abundance of lineages positive for a trait. In (B), values are scaled to the modern value and calculated as (lineages positive for trait x at time t) / (lineages positive for trait x at time 0). A value of 0.5 at 1 Ga means the number of lineages positive for the trait at that time was 50% of the number positive today. See Fig. S2 for values scaled by the historic value. Abbreviations: Ga = billion years ago.

### Predictions were insensitive to most changes in parameters

Our main analysis used a single set of parameters for dating the tree, reconstructing gene functions, and trait prediction. Our next step was to vary these parameters to determine how they affected the outcome of our predictions.

We first varied parameters for dating the tree (Fig. S4). Our main analysis used 10 nodes for calibrating the tree, and in alternate analyses, we removed these calibrations one at a time. In other analyses, we varied the date for the root node, which affected dating of the tree in other studies (*22–25*). As a sanity check, we included an analysis with the tree as dated by Moody et al. (*26*), which involved different calibrations (for geochemical events) and a dating program (McmcTree).

With a few exceptions, the outcome of these alternate analyses for dating was similar to the main analysis (Fig. S4). In most cases, trees in alternate and main analyses appeared similar (Fig. S4A), and node ages displayed a 1:1 relationship (*r* ≈ 1 and CCC ≈ 1) (Fig. S4). We used these trees to date the appearance of traits, and the dates also followed a 1:1 relationship (Fig. S4). One exception was the calibration for the root node (LUCA). Removing the calibration increased node ages by 33% and extended the root to 7.5 Ga (Fig. S4). Because this date is unrealistic, we set an upper bound that was liberal (at 4.65 Ga, or the age of the solar system) or conservative (to 3.448 Ga, or the appearance of stromatolites) (see Supplementary Text). Node ages were <1% higher under the liberal bound and 15% lower under the conservative bound (Fig. S4). The conservative bound is extreme, but if real, our main analysis would overpredict the absolute ages for appearance of traits, with few changes to their order. Despite differences in methodology for dating, the tree from Moody et al. (*26*) was similar to the tree in our main analysis; node ages and dates of trait appearance followed a close relationship (*r* > 0.98 and CCC > 0.94) (Fig. S4). In sum, these analyses show that our predictions are insensitive to most parameters in dating the tree, with the well-known exception of the date of the root.

We next varied methods of reconstructing gene functions (Fig. S5). Our main analysis used the GLD model of Count (*19*), and in alternate analyses, we used the Wagner maximum parsimony model of Count and ALE method used by Moody et al. (*26*). The number of unique gene functions reconstructed was lower for the ALE method (7, 560) than with other methods (9,371). It was also generally lower over evolutionary time within and across genomes (Fig. S5). We used these gene functions to predict lineages positive for traits and dates of their appearance. Overall, the dates for appearance of traits for ALE and GLD methods followed a good relationship (*r* > 0.82, CCC > 0.82) (Fig. S5). The largest disagreement was for a non-metabolic trait (motility). Agreement was even higher for the maximum parsimony method (*r* > 0.88, CCC > 0.88) (Fig. S5). Overall, these analyses suggested that our predictions were reasonably robust to the method for reconstructing gene functions.

Our final analysis involved varying a parameter for enzymes in metabolic networks (Fig. S6). The setting in the main analysis was strict (enzymes must have all subunits), but in an alternate analysis, we used a relaxed setting (all but one subunit can be missing). This pushed back the date of appearance for some traits (Fig. S6). For example, nitrogen fixation appeared at 2.6 Ga in the main analysis (when the last subunit of nitrogenase appeared) and 3.4 Ga in the alternate analysis (when the first subunit appeared). This analysis shows that relaxing parameters for completeness of metabolism results in earlier estimates for its appearance.

Overall, predictions from our main analysis agreed well with alternate analyses. Predictions were most sensitive to the date of the root of the tree, which changed absolute dates of appearance of traits but not their relative chronology.

### Metabolism evolved in three waves

After dating the appearance of traits with our tools, we gathered similar dates from the geochemical record (Supplementary Text and Table S3). We combined these values to construct a timeline for the evolution of metabolism in early microbes (Fig. 4). This suggested that metabolism evolved in three waves.

**Fig. 4.**
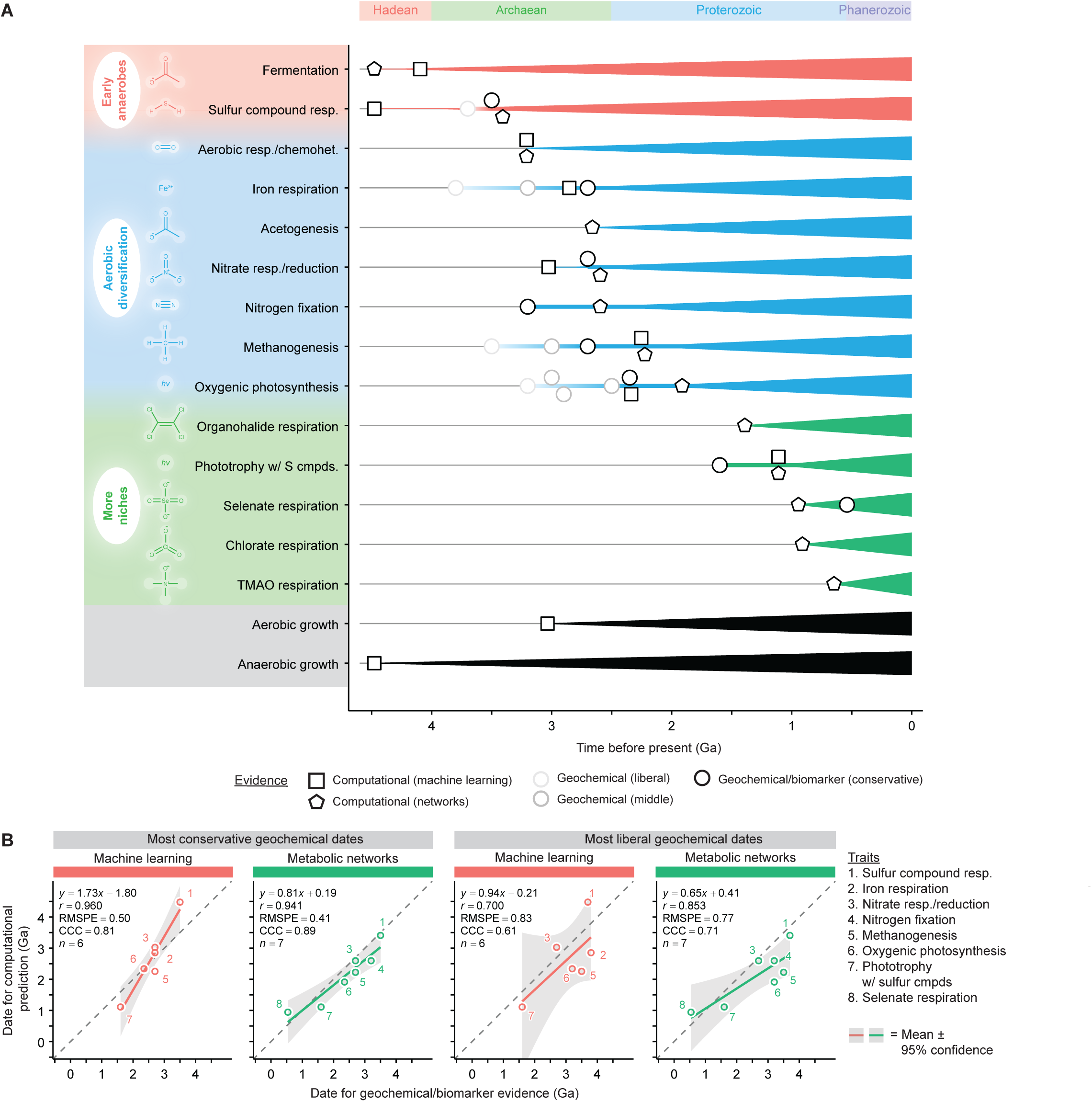
Major forms of metabolism arose in three waves over Earth’s history. See Supplementary Text and Table S3 for dates for geochemical and biomarker evidence. See Table S2 for dates for computational predictions. Abbreviations: CCC = concordance correlation coefficient, chemohet. = chemoheterotrophy, cmpds. = compounds, Ga = billion years ago, *r* = Pearson correlation coefficient, resp. = respiration, RMSPE = root mean square prediction error, TMAO = trimethylamine N-oxide, and w/ = with.

The earliest wave started at ∼3.4 Ga or earlier, and it consisted of early anaerobes carrying out fermentation and sulfur compound respiration (Fig. 4A). These forms of metabolism do not require oxygen, which was absent from early Earth (*31*). Growth was predicted to be strictly anaerobic at this time. The first well-accepted geochemical evidence of any metabolism (for sulfur compound respiration) appears during this time (3.5 Ga) (*32–34*) (Supplementary Text).

The next wave of metabolism started at ∼3.2 Ga and consisted of aerobic diversification (Fig. 4A). It began with aerobic respiration itself, followed by iron respiration, acetogenesis, nitrate respiration, nitrogen fixation, methanogenesis, and oxygenic photosynthesis. We call this aerobic diversification to reflect it began with aerobic respiration (3.2 Ga), aerobic growth (3.0 Ga), and the first evidence of oxygen (3.2 Ga) (*9*) (Supplementary Text). The name does not imply strictly aerobic metabolism, and several forms of metabolism under it do not require oxygen at all. If the most liberal dates for geochemical evidence are accepted, then some of these anaerobic forms of metabolism should be moved to the first wave.

The third wave started at ∼1.4 Ga and involved specialization (Fig. 4A). It consisted of forms of metabolism, such as selenate respiration and chlorate respiration, that are relatively specific and not widely distributed. The last form of metabolism we consider evolved at ∼0.6 Ga. After that, the number of lineages for existing forms of metabolism continued to increase.

When defining these waves, we relied on our computational predictions and the most conservative estimates in the geochemical record. These conservative estimates are the youngest widely accepted minimum age for a given type of metabolism, and the liberal estimates are the oldest and not as widely accepted (Supplementary Text and Table S3). Our computational predictions generally have good agreement with these conservative estimates (Fig. 4B). If liberal values from the geochemical record are included over conservative ones, agreement is lower (Fig. 4B) but still higher than random chance (Fig. S7).

While the waves were named by evolutionary events, their definition was also supported statistically. Silhouette analysis showed the optimal number of waves (k-means clusters) was 3, which is not expected by random chance (Fig. S8). The bounds of the waves include events as shown in Fig. 4.

### Predictions can be explored via a web server

We made our predictions available on Microbe Decoder (https://microbe-decoder.org/?tab=examples) (Fig. S9). With this web server, users can explore predictions on labeled phylogenetic trees. They can see additional details for predictions, such as the specific metabolic pathways predicted by the metabolic networks tool. Users can also make their own predictions (such as with different substrates) by using the reconstructed gene functions included as example data on the server. This allows users to explore our predictions in more detail and test their own hypotheses.

## DISCUSSION

The timeline for the evolution of microbial metabolism is important to Earth’s history, and it is one of the most debated in biology and Earth science (*1–3*). The debate itself dates to a century ago, when Haldane (*35*) claimed that early oceans were a “hot dilute soup” and the first form of metabolism was fermentation. Oparin (*36*) made a similar claim, and they countered the earlier idea that autotrophy came first (*37*, *38*). In years since, the earliest forms of metabolism have been suggested to include sulfur compound respiration (*39*), acetogenesis (*29*, *40*), methanogenesis (*40*, *41*), iron respiration (*42*), and the Wächtershäuser reaction (involving pyrite) (*43*). The evolution of later forms and their abundance over time are even more contentious.

One reason for the debate is that historical evidence for metabolism is ambiguous and patchy. Geochemical evidence of oxygen at 3.2 Ga (*9*) may represent photosynthesis, or it may represent oxygen formed non-biologically from lysis of water by light (*44*) or water reacting with rocks (*45*). Fossil cyanobacteria tie specific organisms to photosynthesis, but the fossil record is ambiguous until 2.0 Ga (*17*) (Supplementary Text and Table S1). Evidence for other forms of metabolism is even patchier.

Our analysis represents a single framework to date major forms of metabolism. It relies on computational reconstruction of gene functions and trait prediction. Many analyses have dated metabolism at the level of individual genes or pathways (*22–26*). Our analysis builds on this by dating whole traits (e.g., fermentation). The predicted dates do not depend on the geochemical record, allowing independent comparison between the two.

Our analysis suggests that metabolism evolved in three waves, starting with fermentation and sulfur compound respiration. It suggests that the second wave started with appearance of aerobic respiration. While aerobic respiration has historically not been dated so early, it aligns with the time that oxygen first appeared (3.2 Ga) (*9*). It also aligns with the appearance of aerobic growth in our study (3.0 Ga), the same trait in other studies (3.2 and 2.7 Ga) (*25*, *46*), and appearance of oxygen-utilizing enzymes in another study (3.1 Ga) (*47*). Our analysis suggests that the final wave involved specialization, with the last major form of metabolism (among those analyzed here) appearing at ∼0.6 Ga.

While our analysis helps fill gaps, there are areas where it can be expanded and improved. It only tracks metabolism carried out by known enzymes, not metabolism carried out non-enzymatically (*48*) or by enzymes that are now extinct. This may explain why our analysis shows no early form of phototrophy, though geochemical evidence for one type (photoferrotrophy) dates to 3.8 Ga (*49*) (Supplementary Text). It dates some traits too late, such as nitrogen fixation (*50*). This has occurred in other analyses (*24*, *51*), and as we found, the date is closer to geochemical evidence if allowing some enzymatic subunits to be absent. It also does not include eukaryotes, which emerged in the fossil record at 1.7 Ga (*52*) and thus contributed to metabolism later in Earth’s history.

Debate over the evolution of metabolism will likely continue over the next century. However, our analysis suggests that different lines of evidence are less disparate than they first appear. The geochemical record includes a range of dates and ambiguities, but our computational predictions with genomes agree well with its most conservative interpretation. Our predictions in turn rely on the fossil record for time calibration. Combining fossils, geochemistry, and genomes provides the clearest picture yet of how microbial metabolism evolved and transformed Earth.

## MATERIALS AND METHODS

### Downloading data

Data for most analyses were downloaded from repositories for Moody et al. (*26*, *29*). The undated phylogenetic tree of 700 genomes and their gene sequences were from a FigShare repository (https://doi.org/10.6084/m9.figshare.24428659). The sequence alignment used to build the tree and fossil calibration file were from a GitHub repository (https://github.com/sabifo4/anoxphoto-divtimes). The phylogenetic tree (rootedtopology2.treefile) was undated, rooted, and followed topology 2 as defined by Moody et al. (*29*). The gene sequences (in the sequences/ directory) were FASTA files named by their gene function (KO ID). Gene functions were annotated by Moody et al. (*29*) using eggNOG-mapper. The sequence alignment (ortho57_aln.phy) was a concatenation of 57 single-copy orthologous genes and contained 8,152 columns (amino acid sites). The fossil calibration file (Calibs_anoxphoto_withArchExclDPANN.txt) contained 10 nodes bounded by 10 dates. These files were used for further steps in our analyses.

Additional data were downloaded from the repositories of Moody et al. (*26*, *29*) and GTDB (*53*). The taxonomy of the 700 genomes of Moody et al. (*29*) was from a FigShare repository (https://doi.org/10.6084/m9.figshare.24428659). Reconstructed gene functions and the dated tree from Moody et al. (*26*) were from a FigShare repository (https://doi.org/10.6084/m9.figshare.27968166). The taxonomy of the GTDB was from their repository (https://data.gtdb.ecogenomic.org/). The taxonomy of the 700 genomes from Moody et al. (*29*) (ArcBacEuk_names.csv) included their names (BinID) and their assigned taxonomy. The taxonomy from the GTDB (bac120_taxonomy_r232.tsv and ar53_taxonomy_r232.tsv) included names (accessions) and assigned taxonomy. Reconstructed gene functions from Moody et al. (*26*) (in the KEGG_genomes_and_anvio_output/ directory) were CSV files named by the genome. The dated tree from Moody et al. (*26*) (FigTree_withLACA_ILN_95HPD.tree) is the version that matches dates in another file in their analysis (dating_results.csv). These files were used for additional analyses.

### Dating the phylogenetic tree

The phylogenetic tree was dated using McmcDate (v. 1.0.0.0) following the general procedure of Davín et al. (*25*). Before using McmcDate, we prepared a treelist using PhyloBayes MPI (v. 1.9) (*54*) and a custom calibration file with an R script.

The input into PhyloBayes MPI was the phylogenetic tree after removing the root. PhyloBayes MPI was run using the LG substitution model (-lg), a single site-profile category (-ncat 1), and four-category discrete-gamma rate heterogeneity (-dgam 4). Two independent chains were run for a minimum of 10,000 iterations. We checked convergence, removed 1,000 iterations from each chain as burn-in, and combined chains into a single TREELIST file. The TREELIST file was used for further analysis.

The input into the R script was the fossil calibration file. The script converted the file to a format readable by McmcDate, then updated values for nodes, dates, and bounds to match Table S1. The final file contained 10 nodes bounded by 10 dates. The value of hard bounds was set to 1⋅10^5^ and soft bounds were 0.025. The custom calibration file was saved as a CSV and used for further analysis.

The inputs into McmcDate were the phylogenetic tree, TREELIST, and custom calibration file. We ran McmcDate using a sparse covariance matrix for approximating likelihood (--likelihood-spec “SparseMultivariateNormal 0.1”) and uncorrelated gamma relaxed clock (--UncorrelatedGamma). It ran for 12,930 iterations and output a .timetree.monitor file. Using that file and an R script, we removed 4,930 iterations as burn-in, then calculated mean age, median age, and 95% highest posterior density (HPD) intervals for every node. The dated tree with mean node ages was saved as a Newick file and used for further analysis.

### Reconstructing gene functions

We reconstructed gene functions for organisms using Count (v. 25.12-beta) (*19*). The inputs into Count were a matrix of gene functions of modern organisms and the phylogenetic tree. We prepared the matrix of gene functions (containing counts of KO IDs by genome) using gene sequences and an R script. We then ran Count using the Gain-loss-duplication (GLD) model, Negative binomial (Pólya) family size distribution, and 4,096 optimization rounds. After obtaining the optimized rate model, we ran Count again to estimate posterior counts of gene functions, which we then output as a CSV file. Using that file and an R script, we summarized gene functions by genome, removing cases with counts < 1. The resulting set of reconstructed gene functions was saved as a CSV file and used for further analysis.

### Trait prediction

We predicted forms of metabolism and other traits using Microbe Decoder (v. 1.0.9) (*27*). The input into Microbe Decoder was the reconstructed gene functions of organisms. We also input the dated tree and taxonomy as metadata (not for prediction). We ran the machine learning tool using default settings and 14 random forest classifiers (models). The output predictions were probabilities of traits. We ran the metabolic networks tool using default settings and 13 network models. The output predictions were metabolic fluxes, which are positive when a pathway exists between a given substrate and end product. By default, tools consider a trait positive when probability > 0.5 or flux > 1. We downloaded CSV files containing the predictions and used them for further analysis.

### Assigning taxonomy

The taxonomy of the 700 genomes from Moody et al. (*26*) is from an older version of GTDB, and we updated it to GTDB v. 232 (*53*). We used a custom R script with the taxonomy from Moody et al. (*26*) and GTDB as inputs. We matched entries by name, and we assigned phyla names manually to any names from Moody et al. (*26*) that did not match.

### Visualization and statistics

We plotted predictions from Microbe Decoder on the dated tree following Fig. S10. We started by mapping predictions onto the tree, with nodes corresponding to organisms and branches corresponding to lineages. Branches were colored if a node (organism) was positive for a trait. Coloring was split at the midpoint if one node was positive and one was negative, which we called the midpoint rule. The tree layout was calculated using R and the ggtree package (*55*) and visualized with the same package.

We counted lineages positive for traits following Fig. S10. By default, we counted lineages the same way as we colored branches of the tree (the midpoint rule). To provide an upper bound for dates, we also used the child rule, where we counted branches (lineages) starting at the child node. To provide a lower bound, we used the parent rule, where we counted starting at the original (parent) node. Counts of positive lineages were then expressed as a fraction of total lineages. Plots were visualized using R and the ggplot2 package (*56*).

Correlation coefficients and related statistics were calculated using base functions of R. Plots were visualized using R and the ggplot2 package (*56*).

### Additional analyses

After completing the main analysis, we varied parameters for dating the tree, reconstructing gene functions, and enzyme completeness.

For analyses with tree dating, we removed one calibration (row) from the custom calibration file, then re-ran McmcDate to get a dated tree. We repeated this for all calibrations, giving 10 dated trees in total. We followed a similar procedure for analyses changing the calibration of the root node (LUCA). The analysis with the dated tree from Moody et al. (*26*) used a custom R script to convert the original TREE file to a Newick. We mapped predictions from Microbe Decoder onto these dated trees.

For analyses with reconstructed gene functions, we used the reconstructed gene functions from Moody et al. (*26*) and converted them to CSV files using a custom R script. We also obtained reconstructed gene functions using Wagner maximum parsimony with Count (*19*) and saved them as CSV files. These CSV files were used as inputs into Microbe Decoder to predict traits, then we mapped predictions onto the dated tree from the main analysis.

For the analysis with enzyme completeness, we predicted traits with Microbe Decoder as above, but we set the “Enzymes must have all subunits” to False. We then mapped predictions onto the dated tree from the main analysis.

## Acknowledgments

### Funding

United States Department of Agriculture National Institute of Food and Agriculture grant 1019985 (T.J.H.)

University of California, Davis Small Grant in Aid of Research (T.J.H.)

### Author contributions

Writing—original draft: T.J.H

Writing—review & editing: T.J.H

### Data, code, and materials availability

All data needed to evaluate the conclusions in the paper are present in the paper and/or the Supplementary Materials. This study did not generate new materials. Code and files related to downloading data, dating the tree, reconstructing gene functions, and assigning taxonomy are available at https://github.com/thackmann/AncientMetabolism and https://doi.org/10.5281/zenodo.21845372. Code for Microbe Decoder is available at https://github.com/thackmann/MicrobeDecoder and https://doi.org/10.5281/zenodo.17553515.

