## Supplementary Text, Figs. S1 to S10, Tables S1 to S3 for "A timeline for evolution of metabolism in early microbes"

Timothy J. Hackmann

**This PDF file includes:**

Supplementary Text
Figs. S1 to S10
Tables S1 to S3

**Other Supplementary Materials for this manuscript include the following:**

Data S1 to S2

### Supplementary Text

#### Choices for fossil calibrations

We chose fossil calibrations for dating the tree according to Table S1 and the rationale below. A major source of the rationale and dates is from ref. (*25*, *26*, *29*, *57–59*). We updated these calibrations with the most-recently characterized evidence and to be as independent from the geochemical record as possible.

*Moon formation (4.52 Ga)*

The moon was formed when a protoplanet (Theia) struck Earth (*60*). This event would have vaporized much of the planet (*61*), killing any life on it. Accordingly, no group could have appeared earlier than this date. Thus, we chose this as an upper bound for all groups, unless a younger event already constrained them. This follows ref. (*25*, *26*, *29*, *57–59*), though the groups are not identical.

The date is from ref. (*62*). It corresponds to the mean date plus the uncertainty provided by the authors. A similar rule applies to other events that serve as an upper bound.

The shape of this bound is hard because evidence is unambiguous and well-dated. Dates older than the bound are effectively not allowed during calibration.

Other events we could have chosen are the formation of the solar system (4.65 Ga) (*63*) or appearance of water (4.40 Ga) (*64*). The formation of the solar system would serve as an upper bound if life started outside Earth.

*Early stromatolites (maximum) (3.448 Ga)*

Stromatolites from the Strelley Pool Formation (*65*) make up the earliest fossil evidence for life that is widely accepted. They have no clearly identifiable cyanobacteria, and claims that they do have alternative interpretations. Following ref. (*26*, *29*, *58*), we chose this as an upper bound for groups containing cyanobacteria (Table S1).

The date is from ref. (*66*) and corresponds to the maximum possible for this formation.

The shape of this bound is soft because cyanobacteria may have existed elsewhere or not been preserved. Dates older than the bound are allowed during calibration but less likely.

Another event we could have chosen is the moon formation (4.52 Ga), but it would be less constraining.

*Early stromatolites (minimum) (3.347 Ga)*

This refers to the same event as above but with the minimum possible date. Thus, life could not have appeared later than this date. Following ref. (*26*, *29*, *57*), we chose this as a lower bound for the last universal common ancestor.

The date is from ref. (*67*) and corresponds to the minimum possible for this formation. It corresponds to the mean date minus the uncertainty provided by the authors. A similar rule applies to other events that serve as a lower bound.

The shape of this bound is hard because evidence is unambiguous and well-dated.

Another event we could have chosen is appearance of carbon of biological origin (4.1 Ga) (*68*). Similarly, we could have chosen the appearance of sulfur compound respiration (3.5 Ga) (*32–34*). However, these events are dated using the geochemical record.

*Methanogens (3.331 Ga)*
Structures from the Kromberg Formation (*69*) are the earliest presumptive fossils of methanogens. Key to this interpretation is their high levels of nickel, a cofactor in methane-forming enzymes. Following ref. (*26*, *59*), we chose this as a lower bound for archaea (excluding DPANN).

The date is from ref. (*70*).

The shape of the bound is soft because microfossils often have other interpretations, and high nickel content is also found in methanotrophs.

Another event we have chosen is appearance of methane of biological origin (*15*), but this event is dated using the geochemical record. The event we ultimately chose still assumes methane existed at this time.

*Stromatolites with gas bubbles (2.720 Ga)*
Stromatolites from the Tumbiana Formation (*71*) have structures interpreted as oxygen bubbles, indicating oxygenic photosynthesis. We chose this as lower bound for total group oxyphotobacteria, which may have been photosynthetic.

The date is from ref. (*72*).

The shape of this bound is soft because the bubbles may have not contained oxygen.

Another event we could have chosen is the first appearance of oxygen (3.2 Ga), but this event is dated using the geochemical record. The event we ultimately chose still assumes oxygen existed at this time.

*Early cyanobacteria (*Eoentophysalis belcherensis*) (2.0136 Ga)*
*Eoentophysalis belcherensis* from the Kasegalik Formation (*17*) is the earliest well-accepted fossil of cyanobacteria. It resembles *Entophysalis*, which belongs to oxyphotobacteria. Following ref. (*25*, *26*, *58*), we chose this as a lower bound for crown group oxyphotobacteria.

The date is from ref. (*73*).

The shape of this bound is soft because this fossil cannot be placed exactly within oxyphotobacteria.

Another event we could have chosen is the widespread appearance of oxygen (2.35 Ga) (*4*, *74–76*), but this event is dated using the geochemical record.

*No eukaryotes (1.8796 Ga)*
Microfossils from the Gunflint Chert are well characterized and do not include eukaryotic cells (*25*, *29*, *58*). Any endosymbionts of eukaryotic cells would be unable to exist in their absence. Following ref. (*77*), we consider classical *Rickettsiales* to be obligate endosymbionts. This group does not include non-classical *Rickettsiales*, which may be free living (*78*). Thus, we chose this event as an upper bound for crown group classical *Rickettsiales*.

The date is from ref. (*79*).

The shape of this bound is soft because eukaryotes may have existed elsewhere.

*Early eukaryotes (1.727 Ga)*
Microfossils from the McDermott Formation (*52*) are the earliest well-accepted evidence of eukaryotes. Presuming *Heimdallarchaeota* were ancestors of eukaryotes (*80*), they would need to have existed beforehand. Thus, we chose this event as a lower bound for crown group *Heimdallarchaeota*. This follows ref. (*26*) but uses a recently-characterized event instead of a younger one (*81*, *82*).

The date is from ref. (*83*).

The shape of this bound is soft because the evidence is not itself a fossil of *Heimdallarchaeota*, but it is instead of a linked group.

Another event we could have chosen is eukaryotes from the Chuanlinggou and Changzhougou Formations (*81*, *82*) (1.6191 Ga), which is the event with the younger date above.

*Thylakoid-bearing cyanobacteria (*Navifusa majensis*) (1.727 Ga)*
*Navifusa majensis* from the McDermott Formation (*18*) is the earliest cyanobacterium that shows clear thylakoids. We chose this event as a lower bound for thylakoid-bearing cyanobacteria, which are oxyphotobacteria excluding *Gloeobacter* and other *Anthocerotibacteraceae*.

The date is from ref. (*83*).

The shape of this bound is hard because evidence is unambiguous and well-dated.

Another event we could have chosen is the appearance of cyanobacteria (*Eoentophysalis belcherensis*) from the Kasegalik Formation (*17*) (2.0136 Ga). This presumes that *Eoentophysalis belcherensis* is indeed a close relative of *Entophysalis* and contained thylakoids.

*Later eukaryotes (*Bangiomorpha pubescens*) (1.030 Ga)*
*Bangiomorpha pubescens* from the Hunting Formation (*84*) is the earliest fossil of a eukaryote belonging to a clear taxonomic group (algae). Any prokaryotic group that donated genes or organelles to eukaryotes would have existed beforehand. The classical *Chlamydiia* donated genes for hydrogenase maturase (*85*), and *Alphaproteobacteria* became mitochondria. Thus, we chose this as a lower bound for total group classical *Chlamydiia* and total group *Alphaproteobacteria*. This follows ref. (*26*), but uses this more certain event over a slightly older one (*86*).

The shape of this bound is soft because the evidence is not itself a fossil of *Chlamydiia* or *Alphaproteobacteria*, but it is instead of a linked group.

The date is from ref. (*87*).

Another event for constraining this group is appearance of *Simia* sp. of the Nonesuch Formation (*86*) (1.08556 Ga), which is the less certain event used by ref. (*26*). Other references (*25*, *29*, *57*, *58*) use *Bangiomorpha pubescens*.

Nostocales *cyanobacteria* (Polysphaeroides filiformis) *(0.993 Ga)*
*Polysphaeroides filiformis* from the Mbuji-Mayi Supergroup (*16*) is the earliest well-accepted fossil of *Nostocales* cyanobacteria. Thus, we chose this event as a lower bound for total group *Nostocales* cyanobacteria. This follows ref. (*25*, *26*) but uses this more certain event over a slightly older one (*88*).

The date is from ref. (*89*).

The shape of this bound is soft because *Polysphaeroides filiformis* do not show heterocysts (*16*), which are present in modern *Nostocales*.

Another event for constraining this group is appearance of *Anhuithrix magna* of the Liulaobei Formation (*88*) (0.996 Ga), which is the less certain event used by ref. (*25*, *26*).

**Choices for geochemical evidence**
We chose geochemical evidence for comparison to computational predictions according to Table S3 and the rationale below. A major source of the rationale is ref. (*2*), with additional sources being ref. (*1*, *3*).

*Sulfur compound respiration*
Sulfur compound respiration has been inferred from samples with low $\delta^{34}S$ and unique values of other sulfur isotopes. In modern microbes, this type of metabolism results in low $\delta^{34}S$ in $H_2S$ (*90*, *91*). A related metabolism, sulfur disproportionation, results in the same (*92*).

The most conservative date for this metabolism is 3.5 Ga (*2*). This corresponds to samples of barite in the Dresser formation with low $\delta^{34}S$ in pyrite (*32–34*). Further, values of $\Delta^{33}S$ and $\Delta^{36}S$ suggest the specific form of metabolism is respiration of sulfate (*32*, *34*) or sulfur disproportionation (*33*), with respiration of sulfur also possible (*33*). This evidence is widely accepted (*2*).

The most liberal date for this metabolism is 3.7 Ga. This corresponds to samples of sedimentary rocks in the Isua Supracrustal Belt with low $\delta^{34}S$ in pyrite (*93*, *94*). Values of $\Delta^{33}S$ and $\Delta^{36}S$ suggest the form of metabolism is respiration of sulfate (*93*) or sulfur (*94*). However, the values are not as extreme as in the Dresser formation.

*Iron respiration*
Iron respiration has been inferred from samples with low $\delta^{56}Fe$, high $\delta^{56}Fe$, or both. In modern microbes, this type of metabolism results in low $\delta^{56}Fe$ in $Fe^{2+}$ and high $\delta^{56}Fe$ in $Fe^{3+}$(*95*).

The most conservative date for this metabolism is 2.7 Ga (*2*). This corresponds to samples of mudstones from the Tumbiana Formation with low $\delta^{56}Fe$ in pyrite (*96*). The authors argue the values of low $\delta^{56}Fe$ are lower than possible through non-biological processes (*97*). The values are even lower than those observed for modern microbes (*95*), but this finding is accepted by others (*2*).

The most liberal date is 3.8 Ga (*2*). This corresponds to samples of iron-rich carbonates from the Isua Supracrustal Belt with high $\delta^{56}Fe$ and low $\delta^{13}C$ (*98*). The authors argue that in their geological setting, the pattern reflects iron respiration. However, it cannot be ruled out that the carbonate originated from younger rocks (*2*).

A middle date is 3.2 Ga (*2*), which corresponds to samples of chert from the Mendon Formation with both low and high $\delta^{56}Fe$ in pyrite (*99*). The authors argue that in their geological setting, this pattern reflects iron respiration. However, they do not rule out non-biological processes, and the values are not as extreme as in the Tumbiana Formation (*2*).

*Nitrate respiration*
Nitrate respiration can be inferred from samples with high $\delta^{15}N$. In modern microbes, this type of metabolism, specifically denitrification (nitrate → $N_2$), results in high $\delta^{15}N$ in nitrate (*100*). It is often preceded by nitrification (ammonium → nitrate), which results in high $\delta^{15}N$ in ammonium (*100*). Any microbial biomass incorporating this nitrate and ammonium would also have high $\delta^{15}N$.

The earliest date for this metabolism is 2.7 Ga (*2*), which corresponds to samples of shale from the Campbellrand-Malmani formation with high values of $\delta^{15}N$ in kerogen (*50*). Other authors have found similar results around this time (*101*).

The earliest date is also a conservative one, with this evidence being widely accepted (*2*).

*Nitrogen fixation*
Nitrogen fixation has been inferred from samples with low $\delta^{15}N$. In modern microbes, this type of metabolism results in low $\delta^{15}N$ in microbial biomass (*102*).

The earliest date is 3.2 Ga (*2*), which corresponds to samples of mudrocks from the Soanesville Group with low values of $\delta^{15}N$ in kerogen (*50*). Similar results were found for other locations slightly younger than this time (*50*).

The earliest date is also a conservative one, with this evidence being widely accepted (*2*).

*Methanogenesis*

Methanogenesis has been inferred from samples with low $\delta^{13}C$. In modern microbes, this type of metabolism results in low $\delta^{13}C$ in methane (*103*) and microbial biomass of methanogens (*104*). If the methane is consumed by other organisms (methanotrophs), $\delta^{13}C$ in microbial biomass would be even lower (*105*).

The most liberal date for this metabolism is 3.5 Ga (*2*). This corresponds to samples of quartz from the Dresser Formation with low values of $\delta^{13}C$ in methane inclusions (*15*). However, non-biological causes cannot be ruled out (*106*). As a result, this evidence is not universally accepted (*2*).

The most conservative date is 2.7 Ga, which corresponds to samples of sedimentary rocks from the Tumbiana Formation with low $\delta^{13}C$ in kerogen (*13*). Other authors have found similar results from this time (*14*). The authors argue the values are so low ($\delta^{13}C \approx$ -60‰) that they originated from methanotrophs consuming biologically-produced methane (*13*). This explanation is widely accepted (*2*), though other forms of metabolism, including acetogenesis (*13*), cannot be ruled out.

A middle date is 3.0 Ga (*2*), which corresponds to samples of shale from the Lalla Rookh Sandstone with low $\delta^{13}C$ in kerogen (*107*). The values are low ($\delta^{13}C \approx$ -35‰), making methanogenesis likely, but they do not rule out other forms of metabolism (*107*).

*Oxygenic photosynthesis*

Oxygenic photosynthesis has been inferred from appearance of oxygen. On modern Earth, this type of metabolism releases $2.8\times10^{17}$ g $O_2$ annually (assuming 1 mol $O_2$ released from 1 mol C of net primary productivity) (*108*). Other biological sources of oxygen include reactions of nitric oxide dismutase, chlorite dismutase, and catalase (*4*). Non-biological sources include lysis of water by light (*44*) or water reacting with rocks (*45*).. The net release of oxygen by these sources is unknown, but on modern Earth, it is accepted to be small compared to oxygenic photosynthesis.

The most conservative date for this metabolism is 2.35 Ga (*4*). This falls within the period when the entire atmosphere was being oxygenated, or the Great Oxidation Event. In this period, $\Delta^{33}S$ of sulfides and sulfates stabilized at 0 (*74–76*), suggesting that photodissociation of $SO_2$ was being stopped by oxygen (ozone). There were also visible changes in sediments. Oxygen-sensitive detrital grains disappeared, while red beds and other oxidized sediments appeared (*4*). These signs together suggest atmospheric oxygen exceeded 0.001% present levels (*4*, *109*). It is universally accepted these levels would require oxygenic photosynthesis.

The most liberal date is 3.2 Ga (*2*), which corresponds to samples of chert from the Manzimnyama Banded Iron Formation having low $\delta^{56}Fe$ and high U (*9*). This has been interpreted as oxygenation of shallow water in that location. However, it is unclear how widespread the oxygenation was and if it could result from non-biological sources.

Even more liberal dates have been suggested (as early as 3.8 Ga) (*12*), but they are not as widely accepted (*2*).

There are several middle dates. Dated at 3.0 Ga, samples of sedimentary rocks from Sinqeni Formation have values of $\delta^{98}Mo$ proportional to Fe/Mn (*10*). At 2.9 Ga, samples of carbonates from the Red Lake greenstone belt have low Ce/Ce*(*11*). At 2.5 Ga, samples of sedimentary rocks from Mount McRae Shale have high Mo and Re (*8*). In all cases, authors interpret these findings as evidence of oxygenation. While this evidence is accepted by many, it has other interpretations. For example, high Mo and Re from Mount McRae Shale has been argued to result from volcanic or postdepositional activity, not oxygenation (*110*).

*Phototrophy with sulfur compounds*

Phototrophy with sulfur compounds has been inferred from samples with the organic molecules chlorobactane, isorenieratane, and okenane. The modern forms of these molecules are chlorobactene, isorenieratene, okenone, and they are found exclusively in purple and green sulfur bacteria (*111*, *112*). All bacteria in these groups carry out this type of metabolism.

The most conservative date is 1.6 Ga. This corresponds to samples from the Barney Creek Formation with chlorobactane, isorenieratane, and okenane (*111*, *112*). Because these markers are found only in these bacteria, this evidence is widely accepted (*2*).

*Selenate respiration*

Selenate respiration has been inferred from low $\delta^{82/78}$Se or similar values for other selenium isotopes. In modern microbes, this metabolism results in low $\delta^{80/76}$Se in selenite and selenium (*113*).

The most conservative date for this metabolism is 0.54 Ga. This corresponds to samples of mudrocks from multiple formations low in $\delta^{82/78}$Se (*114*). Low values can also result from non-biological processes (*115*). However, it is generally accepted that biological processes, such as respiration, are more important (*114*).

*Others*

Other forms of metabolism have been inferred from the geochemical record. However, we do not include them in our analysis because the evidence is ambiguous, the metabolism is not predicted by our computational tools, or both.

Phototrophy with iron (photoferrotrophy) has been inferred from samples with high $\delta^{56}$Fe. In modern microbes, this metabolism oxidizes $Fe^{2+}$ to $Fe^{3+}$, resulting in low $\delta^{56}$Fe in $Fe^{2+}$ and high $\delta^{56}$Fe in $Fe^{3+}$(*98*). The earliest date is 3.8 Ga (*2*), which corresponds to samples of magnetite from the Isua Supracrustal Belt having high $\delta^{56}$Fe (*116*). The authors argue that in their geological setting, this pattern reflects phototrophy with iron. However, high values could also result from another metabolism that oxidizes $Fe^{2+}$(*117*), iron respiration (*98*), or non-biological processes (*118*). We do not include evidence of this metabolism in our analysis because of the ambiguity and because this metabolism is not predicted by our computational tools.

Acetogenesis has been inferred from samples with low $\delta^{13}$C. In modern microbes, this metabolism results in low $\delta^{13}$C in acetate (*119*) and microbial biomass of acetogens (*120*). One early date is 2.7 Ga, which corresponds to samples of sedimentary rocks from the Tumbiana Formation with low $\delta^{13}$C in kerogen (*13*). While these values are accepted as evidence for methanogenesis, another explanation is that they resulted from acetogenesis (*13*). We do not include evidence of this metabolism in our analysis because of the ambiguity.

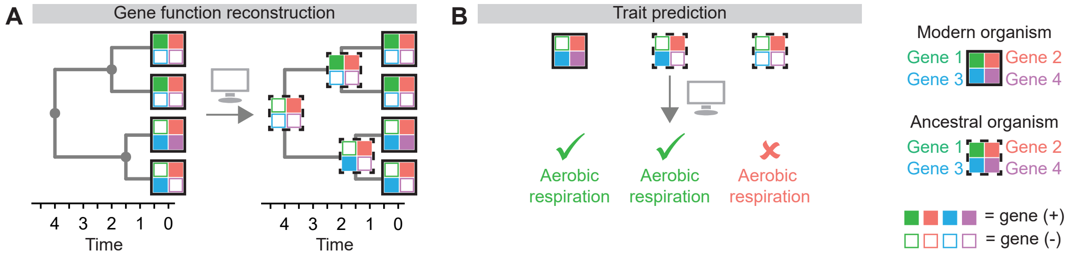


**Fig. S1. Overview of two computational methods used in predicting forms of metabolism in early microbes. (A)** Gene reconstruction. **(B)** Trait prediction.

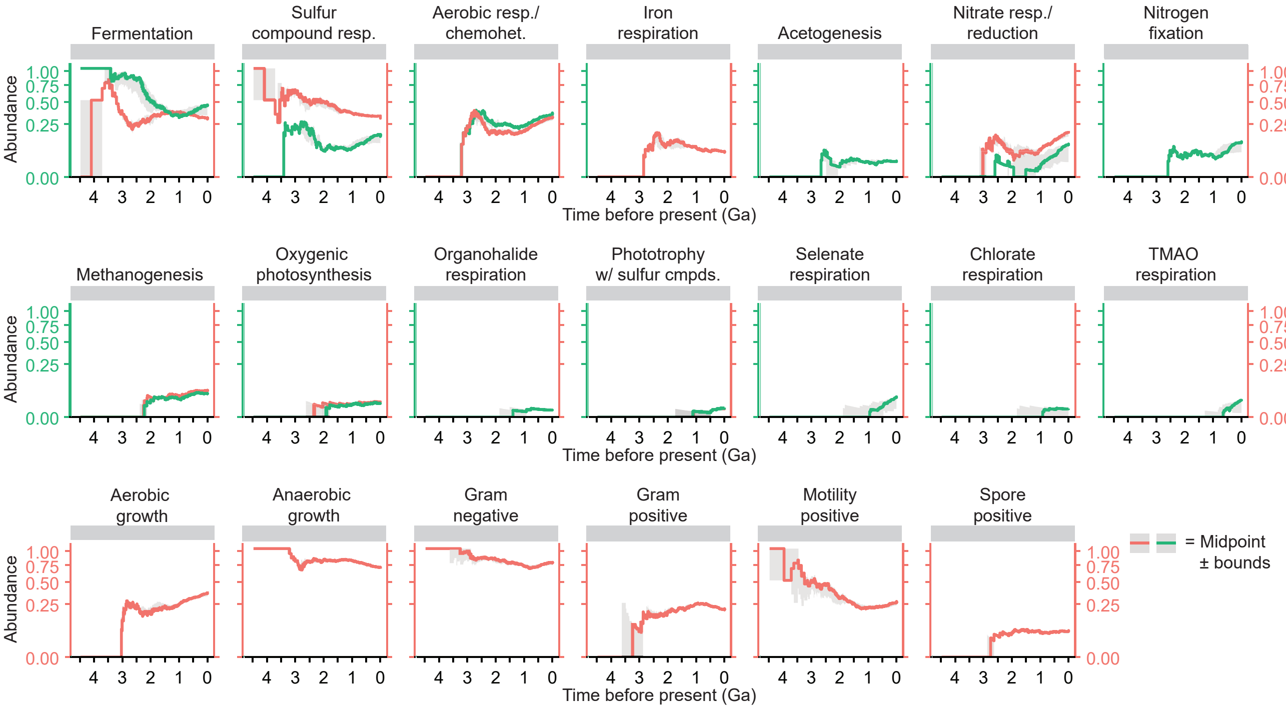


**Fig. S2. Abundance of lineages positive for a trait, scaled to the historic value.** The calculation is (lineages positive for trait *x* at time *t*) / (total lineages at time t). A value of 0.5 at 1 Ga means that 50% of all lineages at 1 Ga were positive for the trait. See Figs. 2 and 3 for values scaled to the modern value. Abbreviations: chemohet. = chemoheterotrophy, cmpds. = compounds, Ga = billion years ago, resp. = respiration, TMAO = trimethylamine N-oxide, and w/ = with.

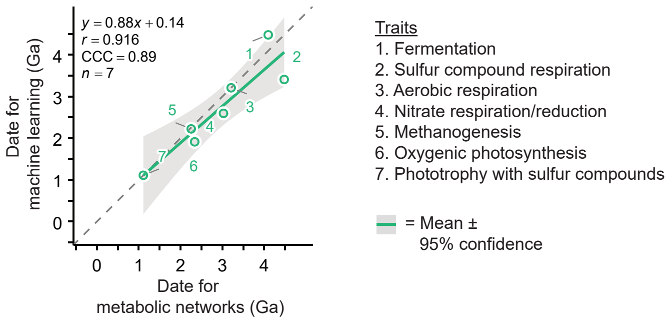


**Fig. S3. Correlation of predicted dates of appearance of traits between tools.** Abbreviations: CCC = concordance correlation coefficient, $r$ = Pearson correlation coefficient, and Ga = billion years ago.

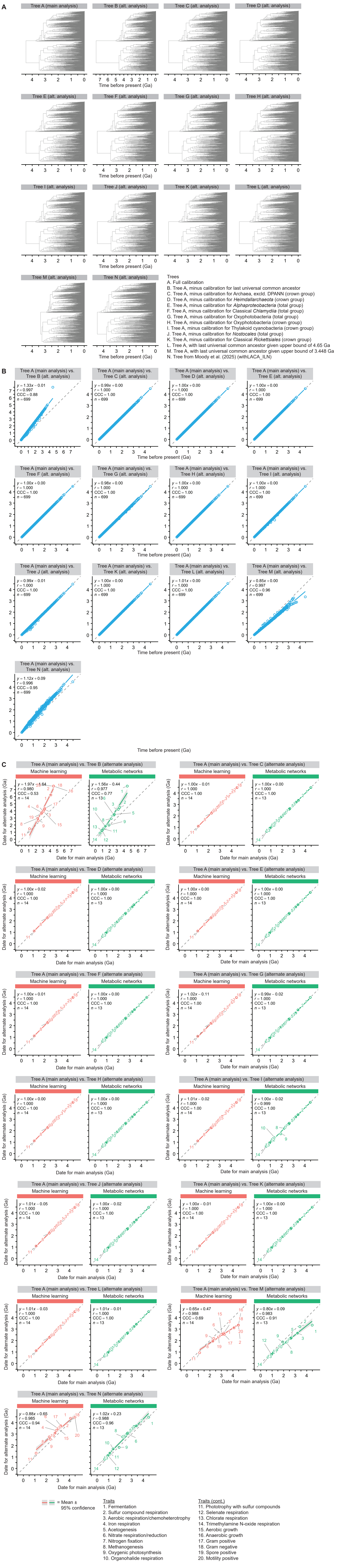


**Fig. S4. Analyses varying parameters for dating the phylogenetic tree. (A)** Trees. **(B)** Correlation between dates for internal nodes. **(C)** Correlation between dates for appearance of traits. In (A) one plot (Tree A) is taken and rescaled from Fig. 1. See Figs. 2 and 3 for results for the main analysis. Abbreviations: CCC = concordance correlation coefficient, $r$ = Pearson correlation coefficient, and Ga = billion years ago.

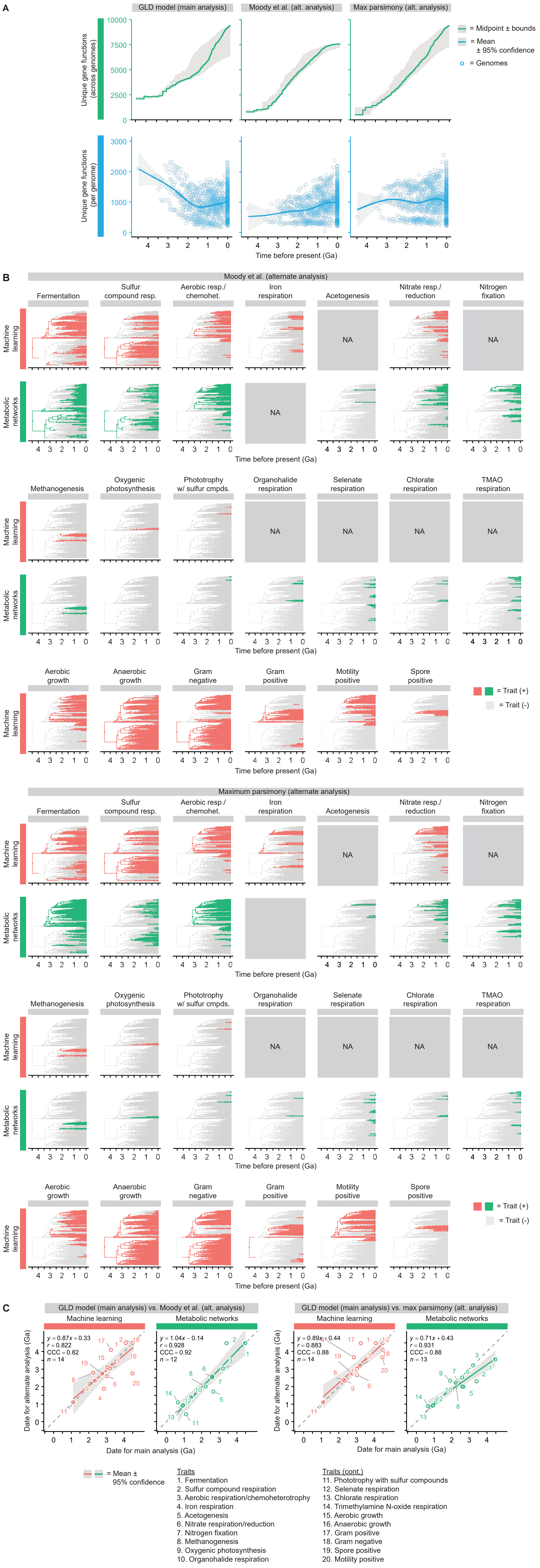

**Fig. S5. Analyses varying methods for reconstructing gene functions.** **(A)** Number of gene functions. **(B)** Lineages positive for traits. **(C)** Correlation between dates for appearance of traits. In (A), one plot (unique gene functions [across genomes] for the GLD model) is taken and rescaled from Fig. 1. See Figs. 2 and 3 for results for the main analysis. Abbreviations: CCC = concordance correlation coefficient, $r$ = Pearson correlation coefficient, and Ga = billion years ago.

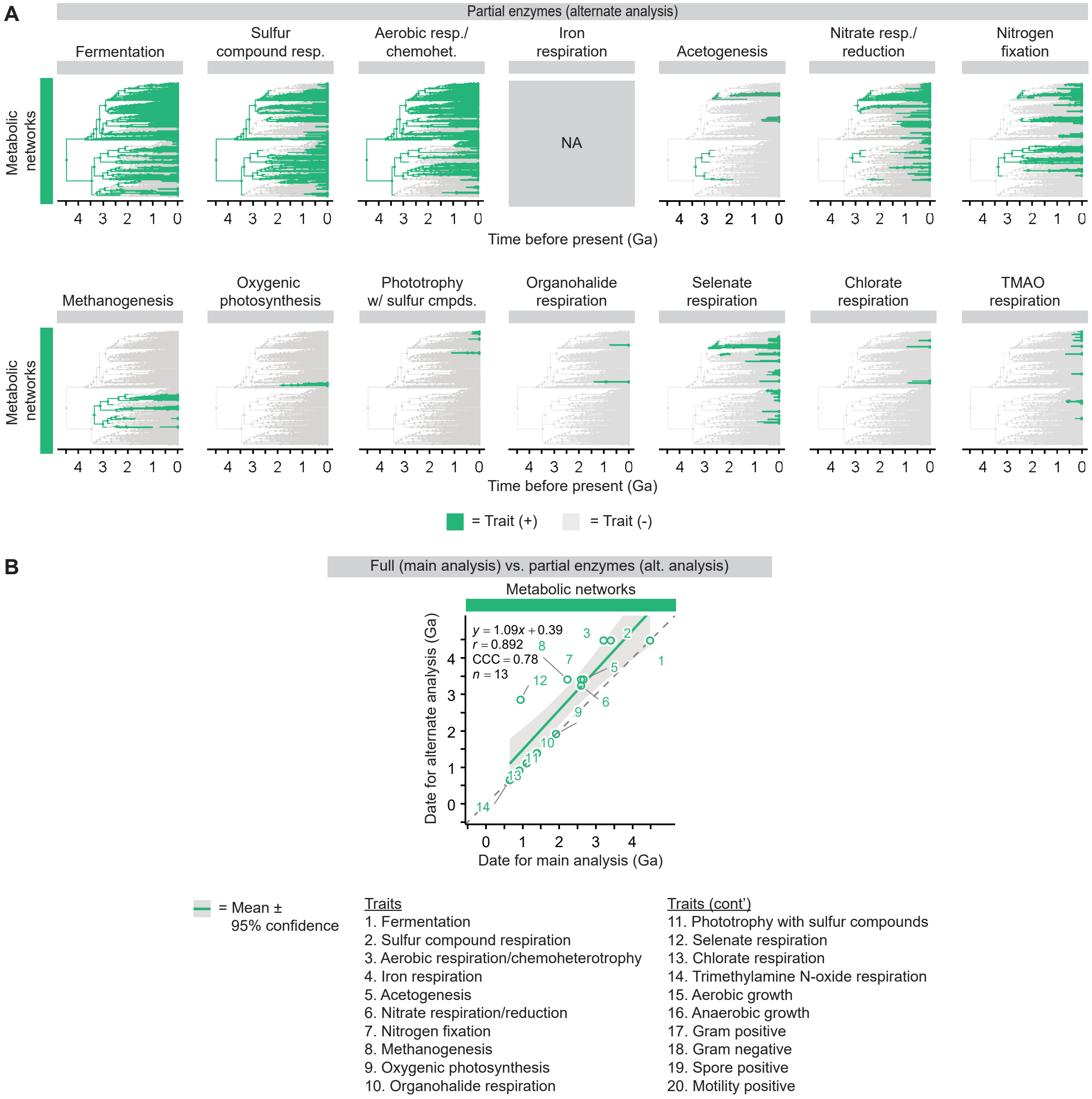


**Fig. S6. Analysis varying a parameter for enzyme completeness. (A)** Lineages positive for traits. **(B)** Correlation between dates for appearance of traits. See Figs. 2 and 3 for results for the main analysis. Abbreviations: CCC = concordance correlation coefficient, *r* = Pearson correlation coefficient, and Ga = billion years ago.

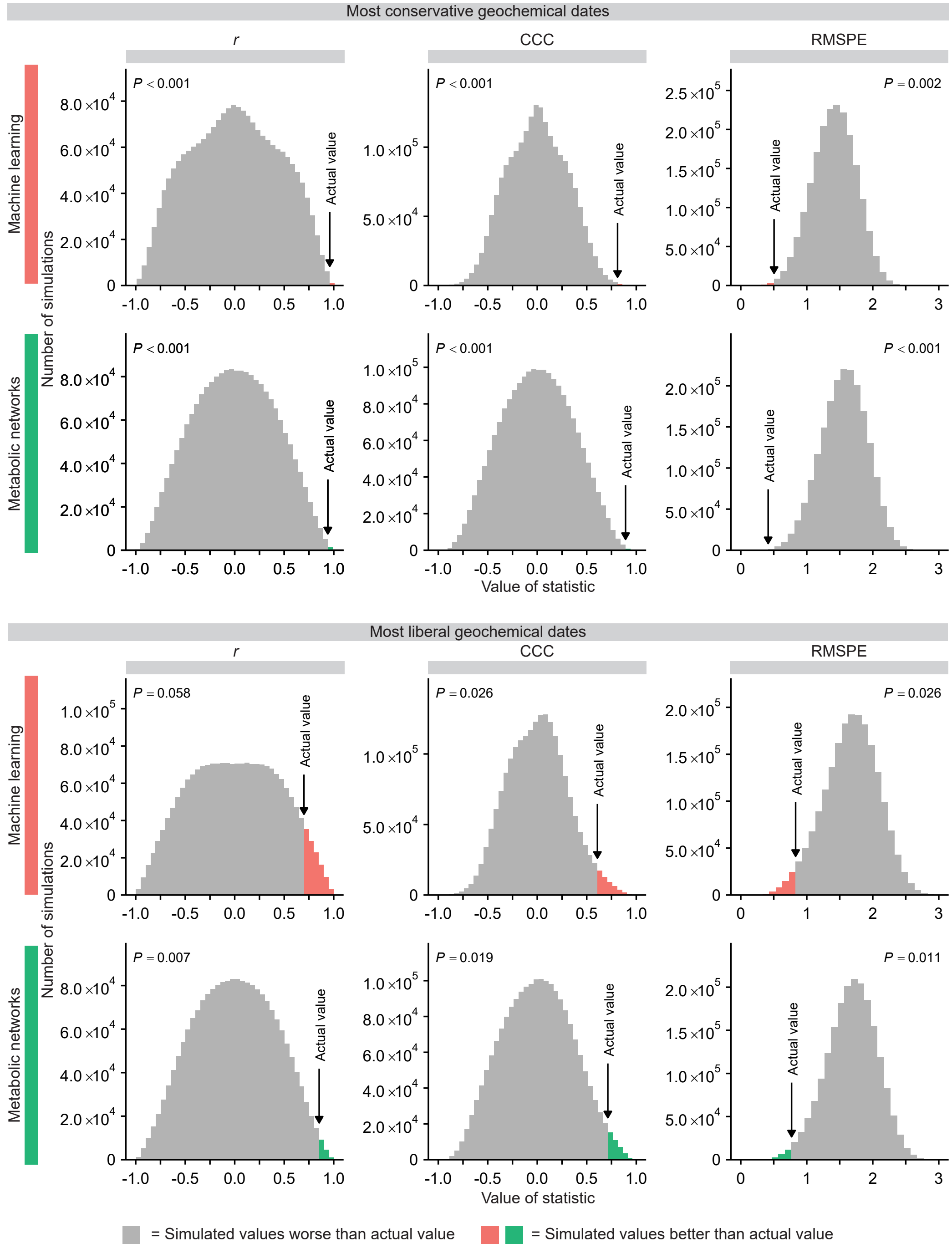


**Fig. S7. Agreement between computational predictions and geochemical evidence is better than random chance, as shown by values of *r*, CCC, and RMSPE from simulations.** In our simulations, we replaced dates from computational predictions with randomly chosen dates (from 0 to 4.48 Ga, the age of the tree root). We then calculated *r*, CCC, and RMSPE between these randomly chosen dates and dates from geochemical evidence. We repeated this process over 2,000,000 simulations. We compared these simulated values of *r*, CCC, and RMSPE to the actual values from the computational predictions (from Fig. 4). The *P*-value is the fraction of simulated values better than the actual values. The low *P*-values show that computational predictions perform better than chance. An exception is *r* for the machine learning tool and most liberal geochemical dates, which has $P > 0.05$. Abbreviations: CCC = concordance correlation coefficient, *r* = Pearson correlation coefficient, and RMSPE = root mean square prediction error.

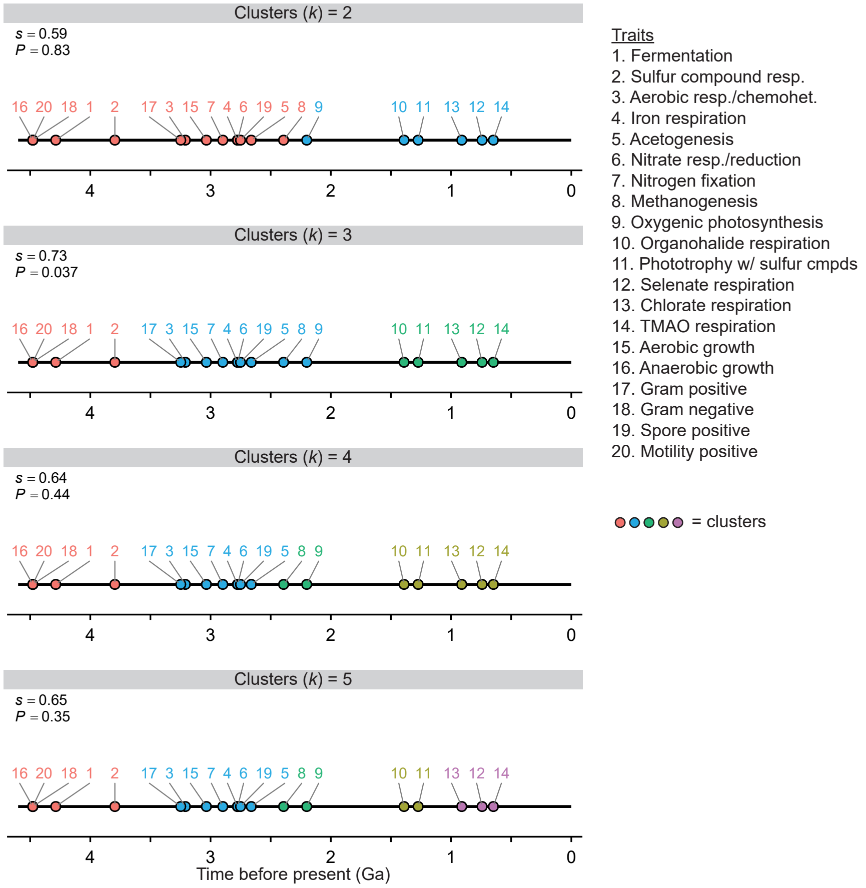


**Fig. S8. Silhouette analysis shows that dates for appearance of metabolism and other traits fall in three clusters.** We took dates for computational predictions (Table S2) and geochemical events (Table S3) and averaged them by trait. The dates for geochemical events were those for the conservative estimate. Missing dates are excluded from the average. We then performed *k*-means clustering on these dates and calculated a silhouette score *s*. We repeated this for $k = 2$ to 5. We then ran Monte Carlo simulations, replacing dates with randomly chosen ones (from 0 to 4.48 Ga, the age of the tree root). We performed *k*-means clustering, calculated *s*, and repeated this process over 10,000 simulations. We compared simulated values of *s* to the actual values above. The *P*-value is the fraction of simulated values better than the actual values. The high value of *s* and low of *P*-value for $k = 3$ indicates that the dates fall into three clusters and that this result is not expected by chance. The bounds of the clusters include events as shown in Fig. 4. Abbreviations: chemohet. = chemoheterotrophy, cmpds. = compounds, Ga = billion years ago, resp. = respiration, TMAO = trimethylamine N-oxide, and w/ = with.

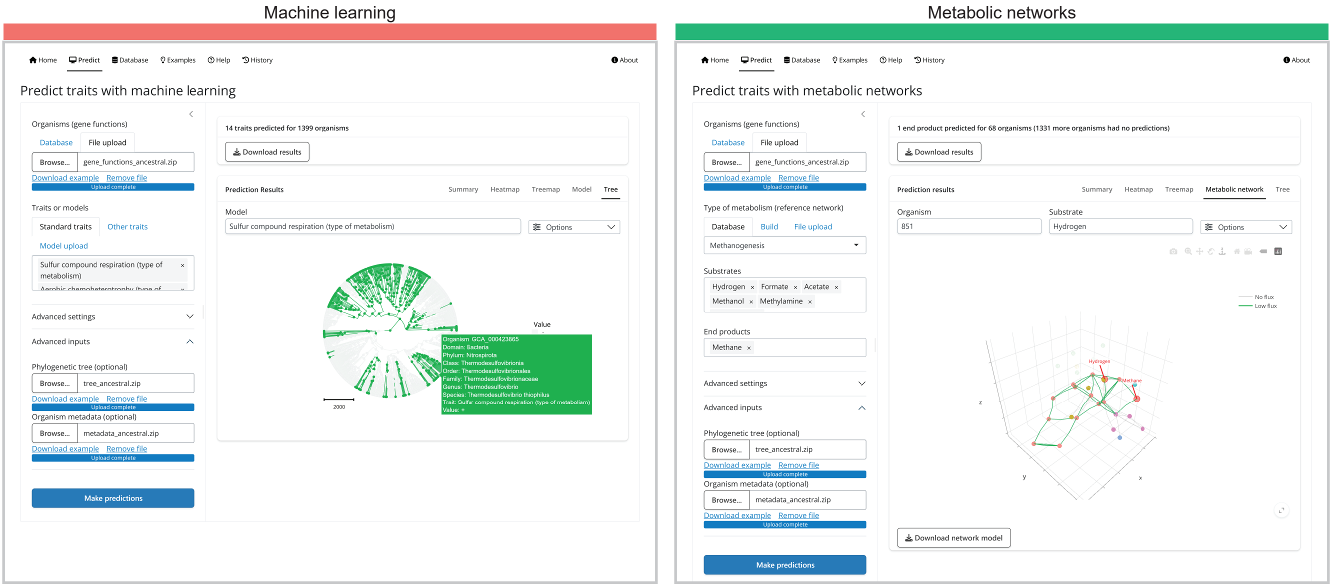


**Fig. S9. Predictions from our analysis can be explored using the Microbe Decoder web server (https://microbe-decoder.org/?tab=examples).**

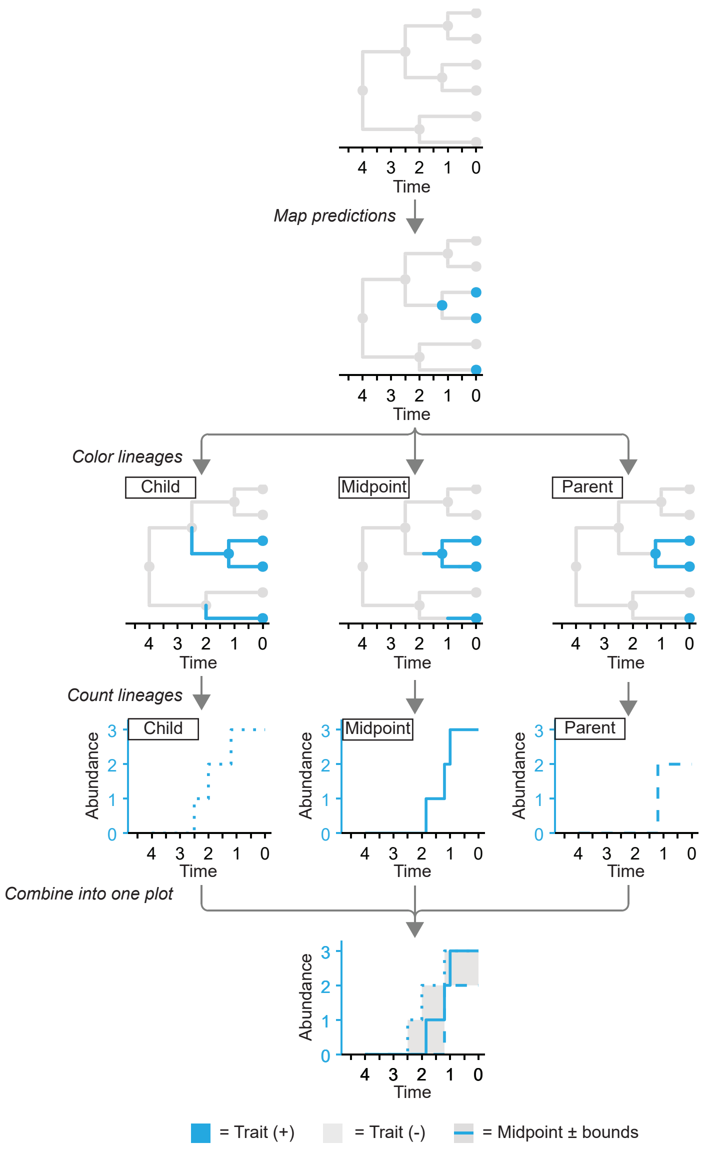


**Fig. S10. Rules for coloring and counting lineages positive for traits in the phylogenetic tree.**

**Table S1. Events used for fossil calibration of the phylogenetic tree.**

| Event | Date (Ga) | Bounds | | | Reference |
|---|---|---|---|---|---|
| | | Group bounded | Direction | Shape | |
| Moon formation | 4.52 | Last universal common ancestor<br>Archaea, excluding DPANN (crown group)<br>*Heimdallarchaeota* (crown group)<br>*Alphaproteobacteria* (total group)<br>Classical *Chlamydiia* (total group)<br>Oxyphotobacteria (total group) | Upper | Hard | (*60*) |
| Early stromatolites (max age) | 3.448 | Oxyphotobacteria (crown group)<br>Thylakoid cyanobacteria (crown group)<br>*Nostocales* (total group) | Upper | Soft | (*65*) |
| Early stromatolites (min age) | 3.347 | Last universal common ancestor | Lower | Hard | (*65*) |
| Methanogens | 3.331 | Archaea, excluding DPANN (crown group) | Lower | Soft | (*69*) |
| Stromatolites with gas bubbles | 2.720 | Oxyphotobacteria (total group) | Lower | Soft | (*71*) |
| Early cyanobacteria (*Eoentophysalis belcherensis*) | 2.0136 | Oxyphotobacteria (crown group) | Lower | Soft | (*17*) |
| No eukaryotes | 1.8796 | Classical *Rickettsiales* (crown group) | Upper | Soft | (*25*, *29*, *58*) |
| Early eukaryotes | 1.727 | *Heimdallarchaeota* (crown group) | Lower | Soft | (*52*) |
| Thylakoid-bearing cyanobacteria (*Navifusa majensis*) | 1.727 | Thylakoid cyanobacteria (crown group) | Lower | Hard | (*18*) |
| Later eukaryotes (*Bangiomorpha pubescens*) | 1.03 | *Alphaproteobacteria* (total group)<br>Classical *Chlamydiia* (total group) | Lower | Soft | (*84*) |
| Nostocales cyanobacteria (*Polysphaeroides filiformis*) | 0.993 | *Nostocales* (total group) | Lower | Soft | (*16*) |

**Table S2. Predicted dates of appearance of traits**

| Tool | Trait | Model | First appearance (Ga) | Oldest bound (Ga) | Youngest bound (Ga) |
|---|---|---|---|---|---|
| Machine learning | Anaerobic growth | Anaerobe (oxygen tolerance) | 4.48 | 4.48 | 4.48 |
| | Gram negative | Gram negative (gram stain) | 4.48 | 4.48 | 4.48 |
| | Motility positive | Motility positive (motility) | 4.48 | 4.48 | 4.48 |
| | Sulfur compound respiration | Sulfur compound respiration (type of metabolism) | 4.48 | 4.48 | 4.48 |
| | Fermentation | Fermentation (type of metabolism) | 4.1 | 4.48 | 3.71 |
| | Gram positive | Gram positive (gram stain) | 3.25 | 3.62 | 2.87 |
| | Aerobic respiration/chemoheterotrophy | Aerobic chemoheterotrophy (type of metabolism) | 3.21 | 3.27 | 3.15 |
| | Aerobic growth | Aerobe (oxygen tolerance) | 3.03 | 3.06 | 3.01 |
| | Nitrate respiration/reduction | Nitrate reduction (type of metabolism) | 3.03 | 3.15 | 2.91 |
| | Iron respiration | Iron respiration (type of metabolism) | 2.85 | 2.91 | 2.8 |
| | Spore positive | Spore positive (spore formation) | 2.75 | 2.87 | 2.63 |
| | Oxygenic photosynthesis | Oxygenic photoautotrophy (type of metabolism) | 2.34 | 2.63 | 2.04 |
| | Methanogenesis | Methanogenesis (type of metabolism) | 2.25 | 2.39 | 2.13 |
| | Phototrophy with sulfur compounds | Anoxygenic photoautotrophy, S oxidizing (type of metabolism) | 1.11 | 1.74 | 0.48 |
| Metabolic networks | Fermentation | Fermentation of glucose | 4.48 | 4.48 | 4.48 |
| | Sulfur compound respiration | Sulfur compound respiration (generic) | 3.41 | 3.46 | 3.36 |
| | Aerobic respiration/chemoheterotrophy | Aerobic respiration (generic) | 3.21 | 3.27 | 3.15 |
| | Acetogenesis | Acetogenesis | 2.66 | 2.69 | 2.63 |
| | Nitrate respiration/reduction | Nitrate respiration (generic) | 2.6 | 2.63 | 2.56 |

| | | | | |
|---|---|---|---|---|
| Nitrogen fixation | Nitrogen fixation | 2.6 | 2.63 | 2.56 |
| Methanogenesis | Methanogenesis | 2.22 | 2.31 | 2.13 |
| Oxygenic photosynthesis | Oxygenic photosynthesis | 1.91 | 2.04 | 1.78 |
| Organohalide respiration | Organohalide respiration (generic) | 1.39 | 1.87 | 0.91 |
| Phototrophy with sulfur compounds | Phototrophy with sulfur compounds | 1.11 | 1.74 | 0.48 |
| Selenate respiration | Selenate respiration (generic) | 0.94 | 1.89 | 0.46 |
| Chlorate respiration | Chlorate respiration (generic) | 0.91 | 1.82 | NA |
| Trimethylamine N-oxide respiration | Trimethylamine N-oxide respiration (generic) | 0.65 | 1.3 | 0.56 |

**Table S3.** Geochemical and biomarker evidence for appearance of major forms of metabolism in microbes.

| Metabolism | Type of estimate | Date (Ga) | Signature | Location | Sample type | Analyzed material | Reference |
|---|---|---|---|---|---|---|---|
| Sulfur compound respiration | Liberal | 3.7 | Low $\delta^{34}S$, with unique $\Delta^{33}S$ or $\Delta^{36}S$ | Isua Supracrustal Belt | Sedimentary rocks (multiple) | Pyrite | (*93*, *94*) |
| | Conservative | 3.5 | Low $\delta^{34}S$, with unique $\Delta^{33}S$ or $\Delta^{36}S$ | Dresser Formation | Barite | Pyrite | (*32–34*) |
| Iron respiration | Liberal | 3.8 | High $\delta^{56}Fe$, low $\delta^{13}C$ | Isua Supracrustal Belt | Iron-rich carbonate | Whole rock | (*98*) |
| | Middle | 3.2 | Low and high $\delta^{56}Fe$ | Mendon Formation | Chert | Pyrite | (*99*) |
| | Conservative | 2.7 | Low $\delta^{56}Fe$ | Tumbiana Formation | Mudstones | Pyrite | (*96*) |
| Nitrate respiration | Conservative | 2.7 | High $\delta^{15}N$ | Campbellrand-Malmani Formation | Shale | Kerogen | (*50*) |
| Nitrogen fixation | Conservative | 3.2 | Low $\delta^{15}N$ | Soanesville Group | Mudrocks | Decarbonated powder, kerogen | (*50*) |
| Methanogenesis | Liberal | 3.5 | Low $\delta^{13}C$ | Dresser Formation | Quartz | Methane inclusions | (*15*) |
| | Middle | 3 | Low $\delta^{13}C$ | Lalla Rookh Sandstone | Shale | Decarbonated powder | (*107*) |
| | Conservative | 2.7 | Low $\delta^{13}C$ | Tumbiana Formation and others | Sedimentary rocks (multiple) | Kerogen | (*13*) |
| Oxygenic photosynthesis | Liberal | 3.2 | Low $\delta^{56}Fe$, high U | Manzimnyama Banded Iron Formation | Chert | Whole rock | (*9*) |
| | Middle | 3 | $\delta^{98}Mo \propto Fe/Mn$ | Sinqeni Formation | Sedimentary rocks (multiple) | Whole rock | (*10*) |
| | Middle | 2.9 | Low Ce/Ce* | Red Lake greenstone belt and others | Carbonates | Whole rock | (*11*) |

| | | | | | | | |
|---|---|---|---|---|---|---|---|
| | Middle | 2.5 | High Mo, high Re | Mount McRae Shale | Sedimentary rocks (multiple) | Whole rock | (*8*) |
| | Conservative | 2.35 | $\Delta^{33}S \approx 0$, others | Multiple | Multiple | Multiple | (*4*, *74–76*) |
| Phototrophy with sulfur compounds | Conservative | 1.6 | Chlorobactane, isorenieratane, okenane | Barney Creek Formation | Siltstones and shales | Bitumen | (*111*, *112*) |
| Selenate respiration | Conservative | 0.54 | Low $\delta^{82/78}Se$ | Multiple | Mudrocks | Whole rock | (*114*) |

**Data S1 (separate file). Predictions from the machine learning tool of Microbe Decoder.**

**Data S2 (separate file). Predictions from the metabolic networks tool of Microbe Decoder.**
